# Novel cytonuclear combinations modify Arabidopsis seed physiology and vigor

**DOI:** 10.1101/370098

**Authors:** Clément Boussardon, Marie-Laure Martin-Magniette, Béatrice Godin, Abdelilah Benamar, Benjamin Vittrant, Sylvie Citerne, Tristan Mary-Huard, David Macherel, Loïc Rajjou, Françoise Budar

**Affiliations:** Institut Jean-Pierre Bourgin, INRA, AgroParisTech, CNRS, Université Paris-Saclay, 78000 Versailles, France; UMR MIA-Paris, AgroParisTech, INRA, Université Paris-Saclay, 75005, Paris, France; Institute of Plant Sciences Paris Saclay IPS2, CNRS, INRA, Université Paris-Sud, Université Evry, Université Paris-Saclay, Bâtiment 630, 91405 Orsay, France; Institute of Plant Sciences Paris-Saclay IPS2, Paris Diderot, Sorbonne Paris-Cité, Bâtiment 630, 91405, Orsay, France; IRHS, Université d’Angers, INRA, Agrocampus Ouest, UMR 1345, SFR 4207 QUASAV, Angers, France; Génétique Quantitative et Evolution–Le Moulon, INRA, Université Paris-Sud, CNRS, AgroParisTech, Université Paris-Saclay, 91190 Gif-sur-Yvette, France

**Keywords:** cytolines, cytonuclear co-adaptation, cytonuclear interaction, dormancy, seed longevity, germination, seed vigor

## Abstract

The influence of intraspecific variation in cytoplasmic genomes and cytonuclear interactions on key seed traits that can impact adaptation and agriculture has not been thoroughly explored, so far. Here, dormancy, germination performance and longevity of seeds have been assessed in Arabidopsis plants with novel cytonuclear combinations that disrupt coadaptation between natural variants of nuclear and cytoplasmic genomes. Although all three traits were affected by cytonuclear reshuffling, the sensitivity of seed traits to cytoplasmic change was dependent on the nuclear background. Both deleterious and, more surprisingly, favorable effects of novel cytonuclear combinations (in comparison with the nuclear parent) were observed, suggesting suboptimal genetic combinations exist in natural populations for these traits. Significant changes on dormancy and germination performance due to specific cytonuclear interacting combinations mainly occurred in opposite directions, in accordance with the previously proposed ‘dormancy continuum’. Consistently, reduced sensitivity to exogenous ABA and faster endogenous ABA decay during germination were observed in a novel cytonuclear combination that also exhibited enhanced longevity and better germination performance, compared to its natural nuclear parent. Cytoplasmic genomes, therefore, represent an additional resource of natural variation for breeding seed vigor traits.

**Issue section:** Growth and development

**Highlight:** Natural variation in Arabidopsis organelles and cytonuclear interactions influence seed dormancy, longevity and germination performance. Enhanced seed vigor was obtained through the creation of novel cytonuclear combinations.

## Introduction

The evolutionary success of flowering plants is largely due to the invention of seeds, which protect and scatter the next generation and provide it with resources upon germination. Germination is a complex process determined by the intrinsic properties of seeds and influenced by biotic and abiotic environment, aging-related damage, which can occur during seed storage, and dormancy, which prevents germination under favorable conditions. In nature, dormancy allows the seed to await for the favorable season for seedling success (Finch-Savage and Footitt, 2017). Once dormancy is released, the capacity of seeds to germinate in a wide range of environmental conditions contributes to adaptation, and germination speed has a fitness impact in competitive situations (Dubois and Cheptou, 2012). Germination is also a trait of primary interest for agriculture, where seed vigor, defined as the properties that ensure a fast and synchronized germination in field conditions (Finch-Savage and Bassel, 2016), is crucial for the quality of seed lots. In contrast to its adaptive role in natural conditions, dormancy is not a desirable trait in crops, as it leads to low and unsynchronized seedling emergence rates in agricultural production (Shu *et al*., 2015). In the laboratory, seed vigor is assessed through the speed and uniformity of germination in favorable conditions, germination performance under stress conditions, and the ability to survive storage (Finch-Savage et al., 2010; Rajjou *et al*., 2012). Seed longevity, i.e. ability to survive and maintain germination performance despite physiological damage occurring during storage, is an important component of seed vigor (Rajjou and Debeaujon, 2008; Sano *et al*., 2016).

*Arabidopsis thaliana* (hereafter Arabidopsis) has been widely used to decipher seed biology traits through mutant studies and genome-wide analyses (North *et al*., 2010). Arabidopsis natural accessions display genetic variation for seed dormancy, longevity and germination vigor (Alonso-Blanco *et al*., 2003;Clerkx *et al*., 2004; Bentsink *et al*., 2010; Vallejo *et al*., 2010; Yuan *et al*., 2016a). For example, the release of seed dormancy during post-harvest (PH) dry storage occurred at variable speeds in 112 Arabidopsis accessions, with the time of storage required to reach 50% of germination ranging from 3.5 to 264 days (Debieu *et al*., 2013). In addition, Arabidopsis natural variation in seed dormancy and longevity has been shown relevant for adaptation in ecological studies (Donohue *et al*., 2005a; Huang *et al*., 2010; Kronholm *et al*., 2012; Debieu *et al*., 2013; Postma *et al*., 2015). A number of quantitative trait loci (QTL) for dormancy, longevity, germination speed, and germination tolerance to stresses have been reported (van Der Schaar *et al*., 1997; Alonso-Blanco *et al*., 2003;Clerkx *et al*., 2004; Galpaz and Reymond, 2010; Bentsink *et al*., 2010; Huang et al., 2010; Joosen *et al*., 2012; Nguyen *et al*., 2012; Yuan *et al*., 2016b). Further major advances in our understanding of the genetic bases of natural variation in seed dormancy and germination are being provided from the functional study of the genes underlying these loci (Bentsink et al., 2006; Shu *et al*., 2015).

Comparatively, little is known about the role of cytoplasmic variation in the establishment of seed traits important for adaptation or agriculture, despite the crucial role of mitochondria and chloroplasts in seed quality and germination performance. Upon imbibition, mitochondrial respiration resumes almost immediately to fuel the very active metabolism which is required during imbibition and germination (Paszkiewicz *et al*., 2017). Plastids are the site of the early steps of ABA and GA syntheses. These two major hormones govern the balance between dormancy and germination through their antagonist effects: ABA promotes the induction and maintenance of seed dormancy, whereas GAs promote germination by stimulating tegument and endosperm rupture and embryo cellular elongation (Finch-Savage and Leubner-Metzger, 2006; Shu *et al*., 2016). Both mitochondria and chloroplasts have endosymbiotic origins and their co-evolution with the nucleus of the host cells has shaped nuclear and organellar genomes, while organizing the compartmentalized metabolism of the plant cell (Dyall *et al*., 2004; Kutschera and Niklas, 2005). The tuning of mitochondrial and chloroplast functions is under a complex and still elusive network of genetic and metabolic regulations (Rurek, 2016; De Souza *et al*., 2017). This is mainly ensured and controlled by nuclear-encoded factors, some of which interact with organelle genes or their products. Numerous genes encoding organellar proteins are expressed during germination and their inactivation often affects the process (Yang *et al*., 2011; Demarsy *et al*., 2011; Savage *et al*., 2013; Sew *et al*., 2016). In addition, transcriptional regulation of specific mitochondrial protein-encoding genes during seed germination has been largely documented (Howell *et al*., 2008; Narsai *et al*., 2011; Law *et al*., 2012). Also, plastid gene expression is required for proper seed development in Arabidopsis (Bryant *et al*., 2011) and maize (Sosso *et al*., 2012). Based on the severe consequences for seed development and germination arising from anomalies in the genetic and physiological activities of plastids and mitochondria, it could be assumed that natural cytoplasmic variation affecting these traits is strongly limited, if not suppressed, by natural selection. However, a few studies have reported results suggesting an impact of natural variation in cytoplasmic genomes on seed physiological traits. A seminal study on reciprocal Arabidopsis F1s and backcross generations showed that cytoplasm variation impacted germination efficiency (Corey *et al*., 1976). A recent report on seed germination performance of reciprocal hybrids between Arabidopsis lyrata populations suggested that a disruption of cytonuclear coadaptation contributed to the lower germination of F1 and F2 hybrid generations (Hämälä *et al*., 2017). Furthermore, in a study using Arabidopsis cytolines, each combining the nuclear genome of a natural variant with the cytoplasmic genomes of a different variant, we previously demonstrated the impact of cytoplasmic natural variation and cytonuclear interactions on adaptive traits in the field, including germination (Roux *et al*., 2016). Here, using the same cytolines, we have investigated further the effect of natural genetic variation within organelles on specific physiological seed traits relevant to adaptation and agriculture, namely dormancy, germination performance and longevity. We found that genetic variation in organelle genomes could impact these traits, depending on the nuclear background. Furthermore, we uncovered novel cytonuclear combinations that displayed higher performance than their natural nuclear parent, suggesting that cytoplasmic variation could potentially be used in breeding of seed traits. The improved germination performance and seed longevity of a novel cytonuclear combination was associated with more efficient endogenous ABA degradation and lower exogenous ABA sensitivity.

## Materials and Methods

### Plant material

Cytolines are genotypes combining the nuclear genome of one parent with the organelle genomes of another conspecific parent. Hereafter, a cytoline possessing the cytoplasm of accession “A” and the nucleus of accession “B” is designated [A]B. We used series of cytolines derived from eight natural accessions of Arabidopsis, namely: Blh-1, Bur-0, Ct-1, Cvi-0, Ita-0, Jea, Oy-0, and Sha, which were selected for their wide genetic diversity (Mckhann *et al*., 2004; Moison *et al*., 2010). Cytolines were obtained by recurrent paternal backcrosses of hybrids from the di-allele cross (Roux *et al*., 2016). Seed stocks of the 64 genotypes (56 cytolines and 8 parental accessions) are available at the Versailles Arabidopsis Stock Center (http://publiclines.versailles.inra.fr/cytoline/index).

Unless specified, the seed stocks used were obtained in a growth chamber and previously used in the study of (Roux et al., 2016). They are conserved in controlled conditions (4°C, 15-30% relative humidity, RH) in the Versailles Arabidopsis Stock Center. These seed lots had been stored for at least 18 months when used for the experiments described here.

Two additional seed productions were realized for dormancy studies in a growth chamber under the same conditions as in Roux *et al*. (2016)(light 16 h at 21°C, dark 8 h at 18°C), using two plants per genotype, each on either of the two available shelves. An additional seed production of the Sha and [Blh-1]Sha lines was realized in the greenhouse, seeds from four individuals were pooled for each genotype.

### Seed dormancy assays

For the 64 genotypes, the depth of dormancy was measured on freshly harvested seeds then at 3, 6 and 9 months of after-ripening (designated 0PH, 3PH, 6PH and 9PH). Seed imbibition was conducted in Petri dishes (Ø 55 mm) on filter papers with three sheets of absorbent paper (Roundfilter paper circles, Ø 45 mm; Schleicher & Schuell, Dassel, Germany) covered by a black membrane filter (ME 25/31, Ø 45 mm; Schleicher & Schuell, Dassel, Germany) wetted with 1.3 mL of ultrapure water. Seeds were then incubated in a controlled culture room under continuous light (Philips TRM HOW/33 RS tubes, 70 μmol.m^−2^.s^−1^) at 15°C or 25°C. Germination (protrusion of the radicle after rupture of the testa and endosperm) was assessed daily during twelve days by visual examination.

Results were analyzed independently for each PH time and considering both germination temperatures in order to better control the variability of germination success, except for 0PH whose only results at 15°C were considered because none of the genotypes germinated at 25°C upon harvest. The [Sha]Cvi-0 and [Ct-1]Jea genotypes were excluded from the analyzed dataset due to missing data for at least one biological replicate.

### Germination performance tests in challenging conditions

The complete set of 64 genotypes was tested for germination performance on water and NaCl, after stratification. Seeds were placed on a blue blotter papers (Anchor Paper Company, SGB1924B) positioned on the germination medium (Phytoblend, 0.5% in water) with or without NaCl, in square Petri dishes (120 x 120 x 17 mm). Because seed germination tolerance to NaCl is variable from one accession to another (Galpaz and Reymond, 2010), each nuclear series was tested on the NaCl concentration (Table S1) that resulted in approximately 50% germination of the parental accession in a preliminary experiment. Each dish contained four to eight genotypes and two or three replicates were realized. Stratification treatment was applied at 4°C in the dark for four days. Then, dishes were transferred into a phytotron system providing continuous light from three sides at 25°C (Sanyo MLR-351H, Versatile environmental test chamber). Germination time was counted from the transfer into the phytotron. For the following of germination kinetics, images were taken at 5 to 6 time points between 24 and 72 hours of germination with a digital camera (Nikon Coolpix-P510, Nikon, France). Germination was visually assessed using ImageJ (http://rsbweb.nih.gov/ij/). The maximum germination percentage (Gmax) was considered reached when no additional germination was observed after another 24 hours.

Following the same procedure, the Sha nuclear series was tested on mannitol (200 mM) or KCl (100 mM). ABA (50 μM) sensitivity tests were identically realized on Sha, [Blh-1]Sha and [Ita-0]Sha, except that germination was monitored visually daily during four days.

### Controlled Deterioration Treatment (CDT)

The CDT was performed in order to mimic natural seed aging (Tesnier and Strookman-Donkers, 2002; Rajjou *et al*., 2008). These experiments were performed on the Sha and Ct-1 nuclear series, on seeds from the Versailles Arabidopsis Stock Center.

Briefly, dry mature seeds were first equilibrated at 75% relative humidity (20°C) during 3 days. After this step, day 0 controls were immediately dried back in a desiccator with Silicagel for three days. Treatment was done by storing the seeds (at 75% RH, using sealed boxes with saturated NaCl solution) for 10, 20, 30 and 40 days at 35°C. After the treatment, seeds were dried back on Silicagel as above. Germination percentage was measured after seed imbibition on water under continuous light at 23°C.

### Measurement of Na and K ion contents in Arabidopsis seeds

A germination experiment was set as described above for germination performance, with or without NaCl (100 mM), except that seeds were collected either immediately after stratification or after 20 hours of germination. Three replicates were realized. Seeds were rinsed three times with 50 mL of sterile water and dried at 100°C overnight. The dry weight of each sample was measured and samples were digested in 2 mL of 70% HNO_3_ for a total of 3h with temperature ramping from 80 to 120°C. Potassium and sodium contents were determined with a Varian AA240FS atomic absorption spectrometer (Agilent Technologies, Santa Clara, Ca, USA) and concentrations calculated by comparison with Na^+^ and K^+^ standards.

### Measurement of ABA content in Arabidopsis seeds

ABA content was measured on dry seeds and on stratified seeds after 6 h of germination, using the same experiment setting as described above for germination performance, with or without NaCl (100 mM). Twenty mg of seeds were ground in a mortar with liquid nitrogen and lyophilized with a freeze-drier. For each sample, 10 mg of freeze-dried powder were extracted with 0.8 mL of acetone/water/acetic acid (80/19/1 v:v:v). ABA stable labelled isotope used as internal standard was prepared as described in Le Roux et al. (2014). Two nanograms of each standard were added to the sample. The extract was vigorously shaken for 1 min, sonicated for 1 min at 25 Hz, shaken for 10 min at 4°C in a Thermomixer (Eppendorf^®^), and then centrifuged (8,000 g, 4°C, 10 min). The supernatants were collected, and the pellets were re-extracted twice with 0.4 mL of the same extraction solution, then vigorously shaken (1 min) and sonicated (1 min; 25 Hz). After the centrifugations, the three supernatants were pooled and dried (final volume of 1.6 mL). Each dry extract was dissolved in 140 μL acetone, filtered, and analyzed using a Waters Acquity ultra performance liquid chromatograph coupled to a Waters Xevo triple quadrupole mass spectrometer TQS (UPLC-ESI-MS/MS). The compounds were separated on a reverse-phase column (Uptisphere C18 UP3HDO, 100*2.1 mm*3μm particle size; Interchim, France) using a flow rate of 0.4 mL.min^−1^ and a binary gradient: (A) acetic acid 0.1 % in water (v/v) and (B) acetonitrile with 0.1 % acetic acid. The following binary gradient was used (t, % A): (0 min, 98 %), (3 min, 70 %), (7.5 min, 50 %), (8.5 min, 5 %), (9.6 min, 0%), (13.2 min, 98 %), (15.7 min, 98 %). Mass spectrometry was conducted in electrospray and Multiple Reaction Monitoring scanning mode (MRM mode), in negative ion mode (see Le Roux *et al*. 2014 for monitored transitions). Relevant instrumental parameters were set as follows: capillary 1.5 kV (negative mode), source block and desolvation gas temperatures 130°C and 500°C, respectively. Nitrogen was used to assist the cone and desolvation (150 L.h^−1^ and 800 L.h^−1^, respectively), argon was used as the collision gas at a flow of 0.18 mL.min^−1^.

### Statistical analyses

All analyses were performed with the R software. For all experiments, the selected models and the contrast tests are detailed in Methods S1.

For all experiments measuring seed germination, germination success was analyzed using a generalized linear model (McCullagh and Nelder, 1989) assuming a binomial distribution with the dispersion parameter fixed at unity and a logit link function.

For each experiment, the model was selected as follows: (i) factors included in the model were defined according to the experimental design; (ii) starting from the most complete model, nested models were sequentially fitted and the best one according to BIC was selected. Once the model selected, the contrasts relevant to the addressed question were tested, and p-values were adjusted using the Bonferroni procedure to control the family-wise error rate (FWER). A contrast was declared significant if its adjusted p-value was lower than 0.05.

We performed a contrast test procedure to detect significant interacting cytoplasm x nucleus combinations: each test concerned a pair of cytoplasms (C1 and C2) and a pair of nuclei (Na and Nb), designed in the text as ‘cytonuclear interacting combination’; a significant p-value indicated that the effect of changing cytoplasm C1 to C2 in the Na nuclear background differed from the effect of the same change in the Nb background, H0 {([C1]Na-[C2]Na) - ([C1]Nb-[C2]Nb) = 0}. The analysis of data from the complete set of 64 genotypes (8 parents and 56 cytolines) thus allows for the testing of 784 cytonuclear interacting combinations (28 pairs of nuclei x 28 pairs of cytoplasms).

ABA content and Na^+^/K^+^ ion ratio were analyzed using linear models. The selection of the model and tests for effects were conducted as described above.

## Results

### Effects of cytoplasmic variations and cytonuclear interactions on seed dormancy

Following on from the observation of differences in germination rate in field-grown Arabidopsis cytolines (Roux *et al*., 2016), we sought to evaluate whether cytoplasmic variation and cytonuclear interactions specifically impacted seed primary dormancy and the kinetics of its release during after ripening storage. Germination of seeds of the complete set of 64 genotypes (8 natural accessions and 56 cytolines) was tested at harvest and three, six and nine months post-harvest (0PH, 3PH, 6PH or 9PH), without any stratification treatment, in two temperature conditions, 15°C and 25°C (Fig. 1 and S1). As expected from the wide variation for dormancy among the panel of parental accessions, the nuclear background had a strong effect on germination. Notably, no or low germination was obtained with seeds of the Ita-0 nuclear series in any conditions. For this reason, these genotypes were excluded from the subsequent analyses. Most seed lots germinated better at 15°C (Fig. 1) than at 25°C (Fig. S1), as previously reported for Col-0 freshly harvested seeds (Leymarie et al., 2012), except those of the Bur-0 nuclear series, which reached almost 100% germination at 25°C at harvest, but displayed lower germination success at 15°C.

**Figure 1.**
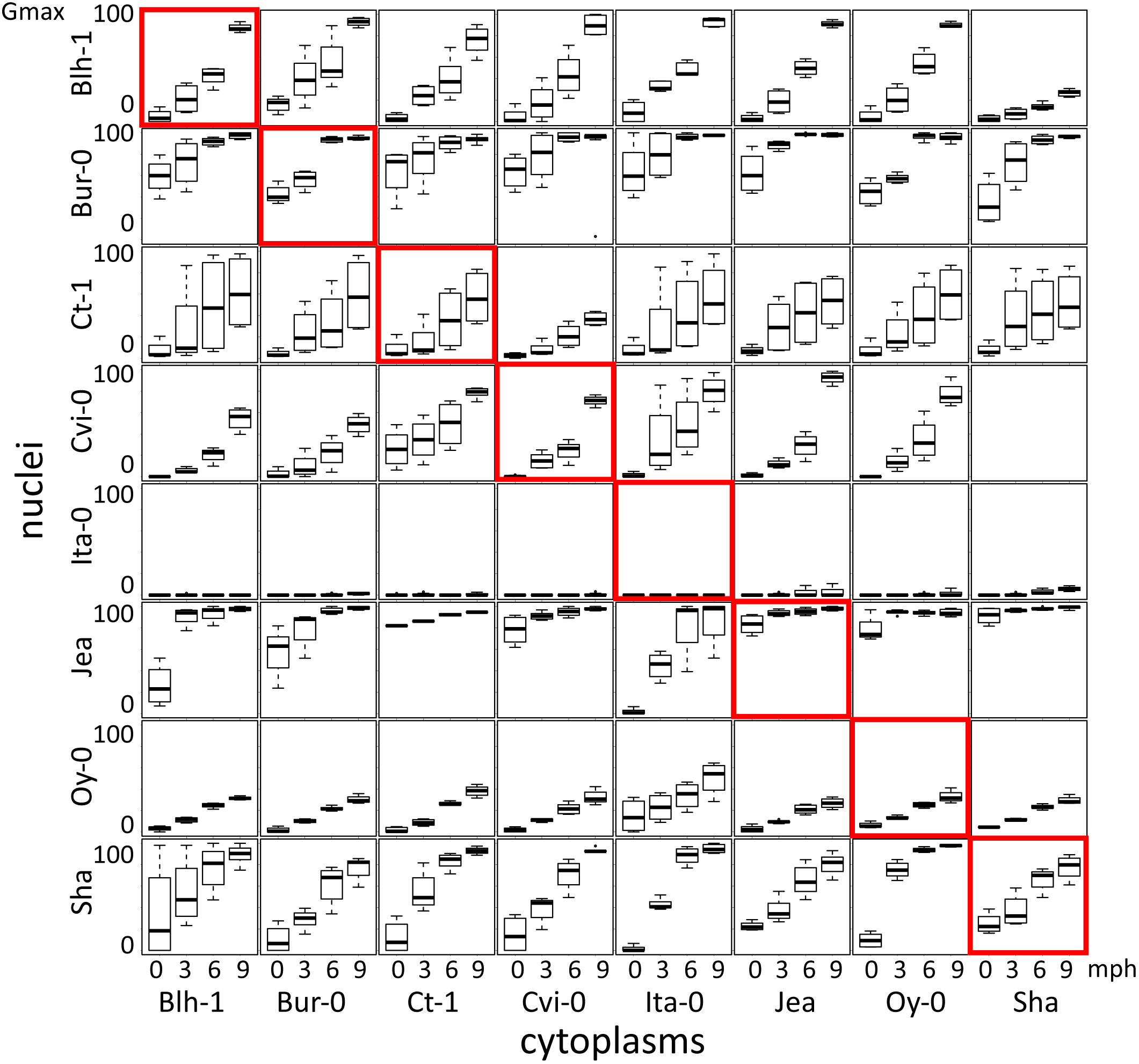
Dormancy release during after ripening in cytoline series. Each panel shows boxplots of the maximum germination percentage at 15°C for one cytonuclear combination 0, 3, 6 and 9 months after harvest (mph). Rows present genotypes sharing their nucleus and columns those sharing their cytoplasm. Natural accessions are framed in red. The [Sha]Cvi-0 cytoline could not be tested due to the small amount of seeds obtained by hand pollination on this male-sterile genotype.

We tested whether germination of cytolines was modified compared to their natural nuclear parent. Table 1 shows that changing organelle genomes had effects on dormancy, but not to the same extent nor in the same direction in all nuclear backgrounds. When effects were observed, foreign cytoplasms tended to enhance dormancy in Ct-1, Jea, Oy-0 and Sha nuclear backgrounds (lower germination in cytolines compared to nuclear natural parent), whereas they rather lowered dormancy in Blh-1, Bur-0, and Cvi-0 backgrounds (higher germination in cytolines). Interestingly, the impact of a given cytoplasm differed according to the nucleus it was associated with, suggesting that dormancy was influenced by cytonuclear interactions. We thus tested for significant cytonuclear interacting combinations impacting dormancy, *i.e*. for differences between two given nuclei in the effect of a given change of cytoplasm, for all possible pairs of cytoplasms and all possible pairs of nuclei (see Material and Methods and Methods S1 for details). As some genotypes were excluded from the analysis due to missing data or total absence of germination, 441 combinations were tested for 0PH data, and 505 for data at other PH times. After Bonferroni corrections for multiple tests, we found 177 significant cytonuclear interacting combinations at 0PH, and 232, 186, and 204 at 3PH, 6PH and 9PH, respectively (Table S2). This indicated that the depth of seed primary dormancy is modulated by genetic variation in interacting nuclear- and organelle-encoded partners.

**Table 1.** Impact of foreign cytoplasms on germination after various post-harvest times in the different nuclear backgrounds.

| Nuclear background<sup>a</sup> | cytoplasm<sup>b</sup> | 0 PH |  | 3 PH |  | 6 PH |  | 9 PH |  |
| --- | --- | --- | --- | --- | --- | --- | --- | --- | --- |
|  |  | Pval<sup>c</sup> | direction of effect<sup>d</sup> | Pval | direction of effect | Pval | direction of effect | Pval | direction of effect |
| Blh-1 | Bur-0 | *** | + | *** | + | *** | + | ns |  |
|  | Ct-1 | ns |  | ns |  | ns |  | * | - |
|  | Cvi-0 | ns |  | ns |  | ns |  | ns |  |
|  | Ita-0 | * | + | ** | + | ns |  | ns |  |
|  | Jea | ns |  | ns |  | ** | + | ns |  |
|  | Oy-0 | ns |  | ns |  | ns |  | ns |  |
|  | Sha | ns |  | ns |  | *** | - | *** | - |
| Bur-0 | Blh-1 | ns |  | ns |  | ns |  | ns |  |
|  | Ct-1 | ** | + | *** | + | ns |  | ns |  |
|  | Cvi-0 | *** | + | * | + | *** | + | ns |  |
|  | Ita-0 | ** | + | *** | + | ns |  | ns |  |
|  | Jea | * | + | *** | + | *** | + | *** | + |
|  | Oy-0 | ns |  | ns |  | ns |  | ns |  |
|  | Sha | ns |  | *** | + | ns |  | * | + |
| Ct-1 | Blh-1 | ns |  | ns |  | ns |  | ns |  |
|  | Bur-0 | ns |  | ns |  | ns |  | ns |  |
|  | Cvi-0 | * | - | ns |  | ** | - | *** | - |
|  | Ita-0 | ns |  | ns |  | ns |  | ns |  |
|  | Jea | ns |  | ns |  | ns |  | ns |  |
|  | Oy-0 | ns |  | ns |  | ns |  | ns |  |
|  | Sha | ns |  | *** | + | ns |  | ns |  |
| Cvi-0 | Blh-1 | NA |  | ns |  | ns |  | ** | - |
|  | Bur-0 | *** | + | ns |  | ns |  | ** | - |
|  | Ct-1 | *** | + | *** | + | *** | + | ns |  |
|  | Ita-0 | * | + | ns |  | *** | + | ns |  |
|  | Jea | NA |  | ns |  | ns |  | *** | + |
|  | Oy-0 | NA |  | ns |  | ns |  | ns |  |
| Jea | Blh-1 | *** | - | ** | - | *** | - | *** | - |
|  | Bur-0 | *** | - | *** | - | ns |  | ns |  |
|  | Cvi-0 | ns |  | * | - | *** | - | *** | - |
|  | Ita-0 | *** | - | *** | - | *** | - | *** | - |
|  | Oy-0 | ns |  | ns |  | ns |  | ns |  |
|  | Sha | ns |  | ns |  | ns |  | ns |  |
| Oy-0 | Blh-1 | ns |  | ns |  | ns |  | ns |  |
|  | Bur-0 | ** | - | ns |  | ns |  | ns |  |
|  | Ct-1 | *** | - | ns |  | ns |  | ns |  |
|  | Cvi-0 | *** | - | ns |  | ns |  | ns |  |
|  | Ita-0 | *** | + | ns |  | ns |  | ns |  |
|  | Jea | * | - | ns |  | ns |  | ns |  |
|  | Sha | ns |  | ns |  | ns |  | ns |  |
| Sha | Blh-1 | ns |  | ns |  | ns |  | * | + |
|  | Bur-0 | *** | - | ns |  | *** | - | ** | - |
|  | Ct-1 | ** | - | ns |  | ns |  | ns |  |
|  | Cvi-0 | ** | - | ns |  | ns |  | ns |  |
|  | Ita-0 | *** | - | ns |  | ns |  | ns |  |
|  | Jea | ns |  | ns |  | ns |  | ns |  |
|  | Oy-0 | *** | - | *** | + | *** | + | * | + |

<sup>a</sup> The results for the Ita-0 nuclear background were not analyzed due to extremely low germination in all conditions (see Fig. 1 and Fig. S1).
<sup>b</sup> For each nuclear background, missing foreign cytoplasms correspond to genotypes excluded before analysis due to missing data or total absence of germination (NAs at OPH).
<sup>c</sup> For each PH time, a difference in germination success between the genotype and its nuclear parent was tested (see Methods S1 for details). For 3PH, 6PH and 9PH experiments, the cytoplasm effects must be interpreted as averaged on germination temperatures. Pvalue : \*\*\*<0.001<\*\*<0.01<\*<0.05; ns, non significant.
<sup>d</sup> the direction of the effect is indicated by the sign of the contrast result. (+) indicates that the tested genotype had a higher germination than its natural nuclear donor. (-) indicates the opposite result.

As seen in Table 1, observed effects of foreign cytoplasms on dormancy often persisted in successive PH times, which was somewhat expected since an effect on the depth of dormancy at harvest would still be detectable after some after-ripening, according to the kinetics of dormancy release. Similarly, we observed persisting effects of cytonuclear interacting combinations over several PH times (Table S2): among the 391 cytonuclear interacting combinations that influenced germination at any PH time, 237 (60%) were significant at two PH times or more and 47 (12%) at all four PH times and, in the vast majority (between 86 % to 100 %) of the cases, they modified germination in the same direction (Table 2), in accordance with a persistence of the observed effects during after-ripening.

**Table 2.** Comparison of directions of the effects of significant cytonuclear interacting combinations shared by two PH times.

| comparison | number of<br>shared<br>cytonuclear<br>interactions | number of<br>identical<br>direction of<br>effects<sup>a</sup> | number of<br>opposite<br>directions of<br>effects<sup>b</sup> |
| --- | --- | --- | --- |
| 0PH vs 3PH | 103 | 95 | 8 |
| 0PH vs 6PH | 82 | 71 | 11 |
| 0PH vs 9PH | 77 | 69 | 8 |
| 3PH vs 6PH | 124 | 123 | 1 |
| 3PH vs 9PH | 110 | 103 | 7 |
| 6PH vs 9PH | 130 | 130 | 0 |

<sup>a</sup> Identical direction of effects corresponds to identical signs of contrast results in Table S2. It indicates that the considered cytonuclear interacting combination affected germination success in the same direction at the two PH times.
<sup>b</sup> Opposite directions of effects corresponds to opposite signs of the contrast results in Table S2. It indicates that the considered cytonuclear interacting combination affected germination success in opposite directions at the two PH times.

Interestingly, the rate of dormancy release during after-ripening also seemed to be modified by foreign cytoplasms in some cases: as seen in Table 1, no difference compared to the natural nuclear parent at 0PH could be followed by either better (*e.g*. [Jea]Cvi-0) or lower (*e.g*. [Sha]Blh-1) germination percentage in later PH times. In addition, a modification in the kinetics of dormancy release was indicated by a change in the contrast sign between PH times in two cases: [Oy-0]Sha germinated less than Sha at 0PH but better at following PH times; oppositely, [Bur-0]Cvi-0 germinated better than Cvi-0 at 0PH, but less at 9PH. Similarly, we interpreted opposite signs of contrast for the effects of a cytonuclear interacting combination significant at to PH times (Table 2) as indication that it altered the kinetics of dormancy release.

### Effects of cytoplasmic variations and cytonuclear interactions on germination performance

In order to assess whether a disruption of cytonuclear co-adaptation also affected the germination performance of seeds after dormancy release, we measured the germination of the complete set of genotypes under permissive and challenging conditions, specifically in the presence of NaCl, at least 18 months after harvest and after stratification. We used salt stress to enhance differences in germination performance that could be difficult to assess after the stratification treatment. Because seed germination tolerance to NaCl is variable from one accession to another (Galpaz and Reymond, 2010), each nuclear series was tested on the NaCl concentration (Table S1) that resulted in 50% germination of the parental accession in a preliminary experiment. Time plots are shown in Fig. S2. Despite the stratification treatment, 100% of germination was not reached on water in Ct-1 and Cvi-0 nuclear series, although germination had reached the plateau (Gmax). Nevertheless, we considered the Gmax as an indicator of germination performance and tested, in each nuclear series, the germination performance of cytolines compared to the natural accession, averaged on the two conditions (Fig. 2, see Methods S1 for details). The results clearly showed that the sensitivity of germination to a cytoplasm change was very variable among the nuclear backgrounds, Blh-1 being very sensitive whereas Ct-1 and Oy-0 were not. In general, when affecting germination, foreign cytoplasms had a negative effect. However, three cytolines ([Ct-1]Ita-0, [Ct-1]Jea and [Blh-1]Sha) did germinate better than their respective nuclear parent. After Bonferroni correction for multiple tests, 202 out of 784 cytonuclear interacting combinations (26%, Table S2) significantly impacted germination success in this experiment. Thus, as dormancy, germination performance was influenced by interactions between organelle- and nuclear-encoded factors that vary among the panel of parental accessions.

**Figure 2.**
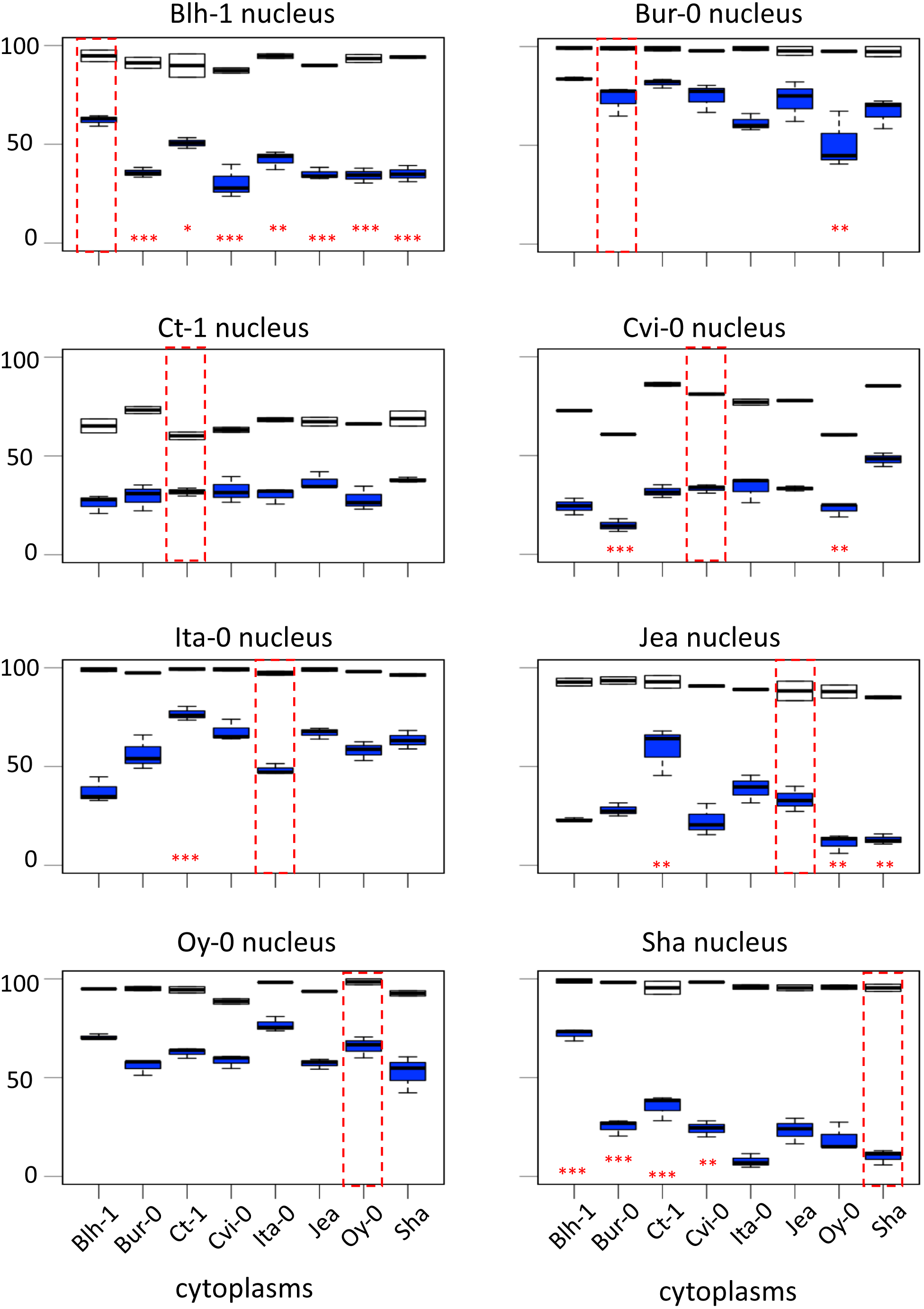
Germination performance of cytoline series. Each panel shows boxplots of the maximum germination percentage (Gmax) of a nuclear series of cytolines, highlighting the natural parental accession in a red dotted frame. White, germination on water; blue, germination on NaCl. Significant differences of germination performance, averaged on the presence of salt, of cytolines compared to their nuclear natural parent are indicated with red stars. Pvalue: ***<0.001<**<0.01<*<0.05 (see Methods S1 for details).

Among the 505 cytonuclear interacting combinations that were tested both for germination performance and for dormancy, 92 were significant for both traits. This represented 80% of the interacting combinations impacting germination performance (92/115), but only 23% of those impacting dormancy for at least one PH (92/392). In addition, the cytonuclear interacting combinations that influenced both dormancy and germination performance impacted germination success more often in the same direction in both experiments than would be randomly expected (Table 3, Chi^2^ test pval <0.001), indicating a negative correlation between dormancy and germination performance.

**Table 3.** Comparison of directions of effects of cytonuclear interacting combinations significant both in the dormancy and germination performance experiments.

| comparison<sup>a</sup> | number of shared significant cytonuclear interacting combinations | number of identical direction of effects<sup>b</sup> | number of opposite directions of effects<sup>c</sup> |
| --- | --- | --- | --- |
| Gmax vs 0PH | 49 | 36 | 13 |
| Gmax vs 3PH | 59 | 45 | 14 |
| Gmax vs 6PH | 37 | 20 | 17 |
| Gmax vs 9PH | 43 | 30 | 13 |

<sup>a</sup> Gmax stands for germination success in the experiment testing germination performance; xPH stands for germination success after x months PH in the dormancy experiment.
<sup>b</sup> Identical direction of the effects correspond to identical signs of the contrasts results in Table S2. Based on the writing of the contrasts, it indicates that the considered cytonuclear interacting combination affected germination success in the same direction in both experiments, thus dormancy and germination performance in opposite directions.
<sup>c</sup> On the same basis, opposite directions of the effects correspond to opposite signs of the contrasts results in Table S2. It indicates that the considered cytonuclear interacting combination affected germination success in opposite directions in both experiments, thus dormancy and germination performance in the same direction.

### Contribution of individual accessions to cytonuclear interactions that impact germination

The above analyses, the impact on germination traits of a shuffling of genetic compartments depended on the origin of the genomes. To better assess this variation, we analyzed the contribution of individual accessions to cytonuclear interacting combinations affecting dormancy or germination performance. Fig. 3 indicates, for each accession, the number of significant cytonuclear interacting combinations that involved its cytoplasm or its nucleus and that impacted dormancy depth, conservatively estimated as effects maintained through all PH times (Fig. 3a) or germination performance after storage (Fig. 3b). Among the 47 cytonuclear interacting combinations that impacted dormancy depth, 28 involved the Ita-0 cytoplasm and 33 involved the Jea nucleus. Among the 202 significant interacting combinations that affected germination performance after storage, 100 involved the Blh-1 cytoplasm and 76 the Sha nucleus. When considering the contribution of accessions to cytonuclear interacting combinations that affect both dormancy and germination performance, the Blh-1 and Ita-0 cytoplasms and the Sha nucleus clearly emerged (Fig.S3).

**Figure 3.**
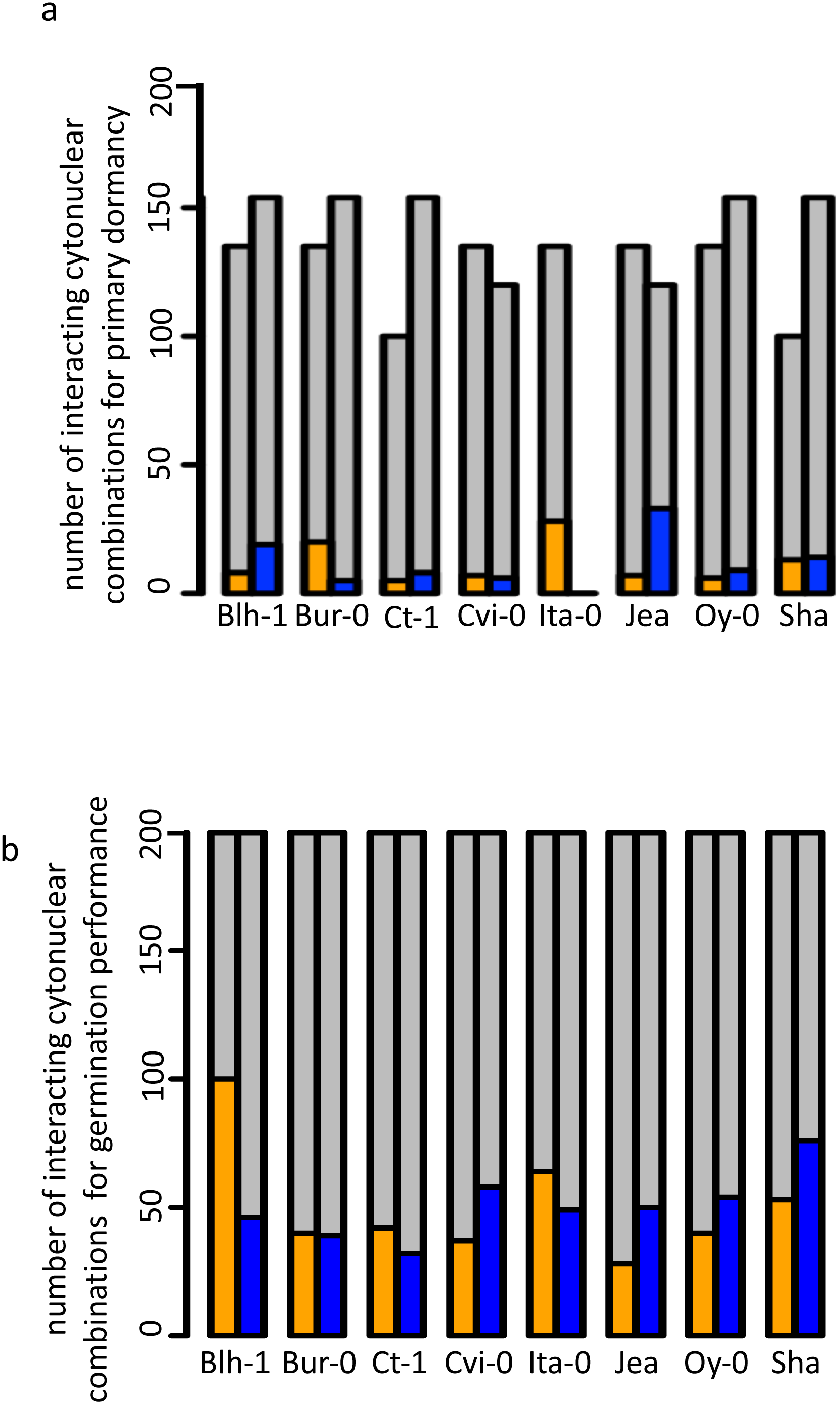
Contribution of individual accessions to cytonuclear interacting combinations impacting germination traits. Cytonuclear interacting combinations are plotted according to the contribution of individual accessions. For each accession, the number of combinations involving its cytoplasm (orange) or its nucleus (blue) that significantly impacted dormancy depth at the four PH times (panel a) or germination performance after storage (panel b) are indicated upon the total number of tested cytonuclear interacting combinations (grey).

### Effects of cytoplasm variation on seed longevity in Sha and Ct-1 nuclear backgrounds

Along with dormancy and germination performance, longevity is an important component of seed vigor. Thus, we addressed whether this trait was also under the influence of cytoplasmic variation. We selected the Sha and Ct-1 series because these two nuclear backgrounds represented two extreme cases in their contribution to cytonuclear effects in the previous experiments (Fig. 3 and S3). The longevity of seeds from these two cytoline series was estimated by measuring their viability (germination percentage in standard conditions) after 0, 10, 20, 30 or 40 days of controlled deterioration treatment (CDT) (Fig. 4). We used the same seed lots as in the assays for germination performance after storage. In this experiment, all genotypes of the Ct-1 series reached 100% germination. We concluded that the partial seed imbibition followed by desiccation included in the CDT treatment, even for the control, acted as a priming treatment releasing the residual dormancy observed in the experiment on germination performance described above. The kinetics of viability loss were different in the two nuclear backgrounds, revealing an expected nuclear effect on seed longevity (Fig. 4). Interestingly, four cytolines appeared less sensitive to CDT than their natural nuclear parent, whereas none was more sensitive (Table 4, see Methods S1 for details). In addition, in both nuclear backgrounds, the selected statistical models included an interaction between the cytoplasm and the length of CDT treatment, which suggested that the loss of germination ability between two treatment durations was under the influence of the cytoplasmic genomes. We could not test directly for significant cytonuclear interacting combinations, because the two series were measured in different experiments and the results analyzed independently. Nevertheless, in Blh-1/Sha and Ita-0/Sha cytoplasm pairs, individual cytoplasms had opposite effects in the two nuclear backgrounds (Table S3), indicating an influence of cytonuclear epistasis on germination after CDT. Strikingly, the Sha cytoplasm performed better than the Blh-1 and the Ita-0 cytoplasms in the Ct-1 nuclear background but worse when associated with its natural nuclear partner.

**Figure 4.**
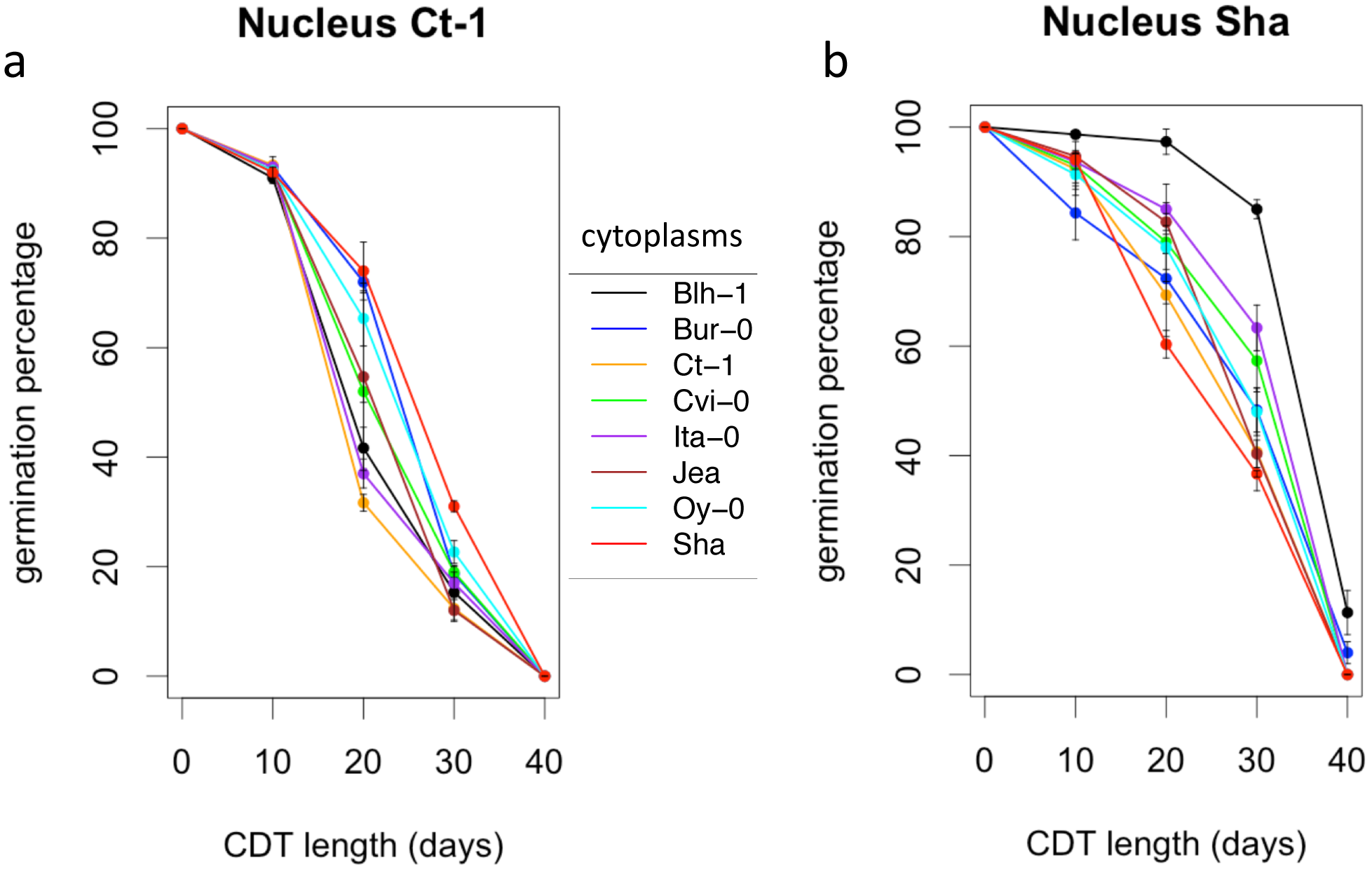
Kinetics of loss of germination efficiency during artificial aging. Controlled deterioration treatment (35°C, 75% relative humidity) was applied for different durations to seeds from cytolines of the Ct-1 (a) and Sha (b) nuclear series. The means of maximum germination percentage from three replicates were plotted. Vertical bars indicate standard deviations from the mean. The colors of the graphs indicate the cytoplasms of the genotypes: black, Blh-1; blue, Bur-0; orange, Ct-1; green, Cvi-0; purple, Ita-0; brown, Jea; cyan, Oy-0; red, Sha.

**Table 4.** Impact of foreign cytoplasms on germination after CDT in the Sha and Ct-1 nuclear backgrounds

| cytoplasm<sup>a</sup> | Ct-1 nuclear background |  | Sha nuclear background |  |
| --- | --- | --- | --- | --- |
|  | Pval<sup>b</sup> | direction of effect<sup>c</sup> | Pval | direction of effect |
| Blh-1 | ns |  | *** | + |
| Bur-0 | * | + | ns |  |
| Ct-1 |  |  | ns |  |
| Cvi-0 | ns |  | ns |  |
| Ita-0 | ns |  | * | + |
| Jea | ns |  | ns |  |
| Oy-0 | ns |  | ns |  |
| Sha | *** | + |  |  |

<sup>a</sup> Each foreign cytoplasm was tested for effect compared to the parental cytoplasm (see Methods S1 for details). The cytoplasm effects must be interpreted as averaged on {10, 20 ,30} days of CDT.
<sup>b</sup> Pvalue : \*\*\*<0.001<\*\*<0.01<\*<0.05; ns, non significant.
<sup>c</sup> (+) indicates that the tested genotype had a higher germination after CDT than its natural nuclear donor. (-) indicates that the tested genotype had a lower germination after CDT than its natural nuclear donor.

### Physiological clues for germination performance of the [Blh-1]Sha line

A closer look at the 202 cytonuclear interacting combinations that influence germination performance revealed that the [Blh-1]Sha combination was involved in 46 (23%, Table S2) of them, indicating that this cytonuclear combination had a major effect on germination. In addition, this genetic combination also had a strong impact on CDT tolerance. We thus decided to get further insight into the behavior of [Blh-1]Sha seeds.

In order to test whether the peculiar behavior of the [Blh-1]Sha seeds was due to the conditions of seed production, we tested germination performance of [Blh-1]Sha and Sha seeds from two productions in different environments, growth chamber and greenhouse. We observed higher germination performance of the cytoline compared to the natural accession for both seed productions (Fig. S4). An environmental cause of the higher germination performance of [Blh-1]Sha was therefore very unlikely. However, we also observed that this positive effect did not persist in seed samples that had been stored several months at room temperature in the laboratory.

We then tested whether the germination performance of [Blh-1]Sha was possibly maintained in the presence of mannitol (200 mM) and KCl (100 mM), by analyzing the germination efficiency of the Sha cytoline series (Fig. S5). The concentrations applied were chosen to compare results to those of the NaCl treatment (100 mM) in the germination performance experiment. The germination rate of the [Blh-1]Sha cytoline was consistently higher than that of the other genotypes under both osmotic and ionic stresses, although these stresses had a milder effect than the NaCl treatment on all genotypes, as previously reported (Munns and Tester, 2008). The germination curve of [Blh-1]Sha on KCl grouped with those of the other lines in the absence of KCl.

It was conceivable that the better germination performance of [Blh-1]Sha seeds would be amplified by an increased tolerance to salt in germination under NaCl stress. Since Na^+^/K^+^ ratio is a key indicator of salt tolerance (Shabala and Cuin, 2008), potassium and sodium contents were analyzed in seeds of [Blh-1]Sha, [Ita-0]Sha and Sha. The [Ita-0]Sha was used as a control cytoline with germination performance similar to Sha (Fig. 2). Very little sodium accumulated in seeds in the absence of NaCl. When germinated on NaCl, both [Blh-1]Sha and [Ita-0]Sha seeds showed a lower Na^+^/K^+^ ratio than Sha seeds (Fig. 5). As the two cytolines had different germination responses on NaCl compared to Sha, we considered it unlikely that the Na^+^/K^+^ balance is the primary cause of the increased germination success of the [Blh-1]Sha cytoline on NaCl. These results reinforced our hypothesis that the intrinsic germination capacity of [Blh-1]Sha seeds was enhanced compared to its natural parent.

**Figure 5.**
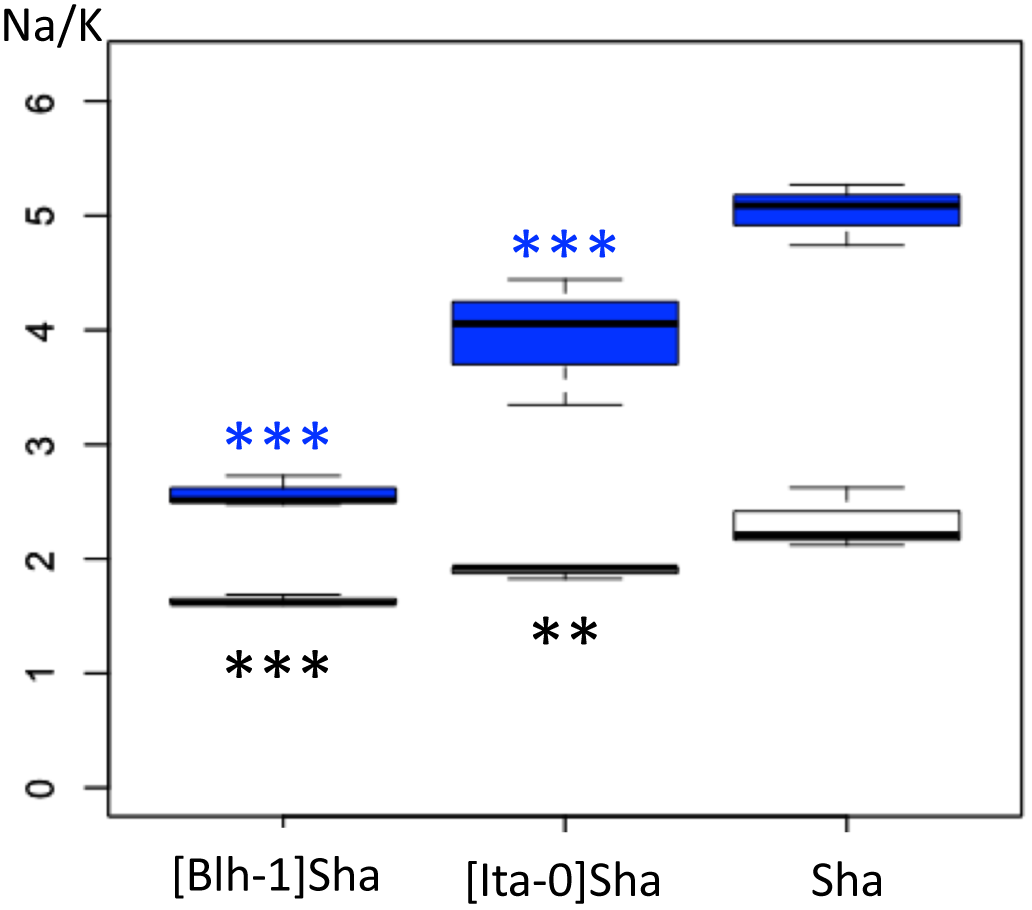
Ratios of Na+ and K+ ion contents in seeds from lines with different cytoplasms associated with the Sha nucleus Ratios of Na+ and K+ ion contents just after stratification (white boxplots) or after 20h of germination (blue boxplots) measured in seeds on 100 mM NaCl. The Na/K ratios were different between the three genotypes in pairwise contrast tests. On the figure, only p-values for comparisons of the cytolines to Sha are shown, ***<0.001<**<0.01.

We then focused on ABA content and sensitivity to further explore physiological parameters that could contribute to the enhanced germination performance of [Blh-1]Sha seeds. We measured the endogenous ABA content of seeds from [Blh-1]Sha, [Ita-0]Sha, and Sha lines. ABA contents were measured in both dry and stratified seeds placed under germination conditions for six hours, in the presence or absence of NaCl (Fig. 6a). The experiment was repeated after 10 further months of storage in controlled conditions (Versailles Stock Centre), yielding similar results. The combined analysis of data from both experiments (see Methods S1 for details) indicated that dry seeds of [Blh-1]Sha contained more ABA than those of [Ita-0]Sha and Sha (p-values < 0.001), whereas they contained less ABA after six hours of germination, either on water or NaCl (p-values < 0.01). The sensitivity of germinating seeds to exogenous ABA (50 μM) was tested on the same seed stocks after the second experiment (Fig. 6b). The [Blh-1]Sha seeds were less sensitive to ABA treatment than Sha seeds. Taken together, the results suggest that both endogenous ABA metabolism and ABA sensitivity are modified in [Blh-1]Sha seeds compared to Sha seeds.

**Figure 6.**
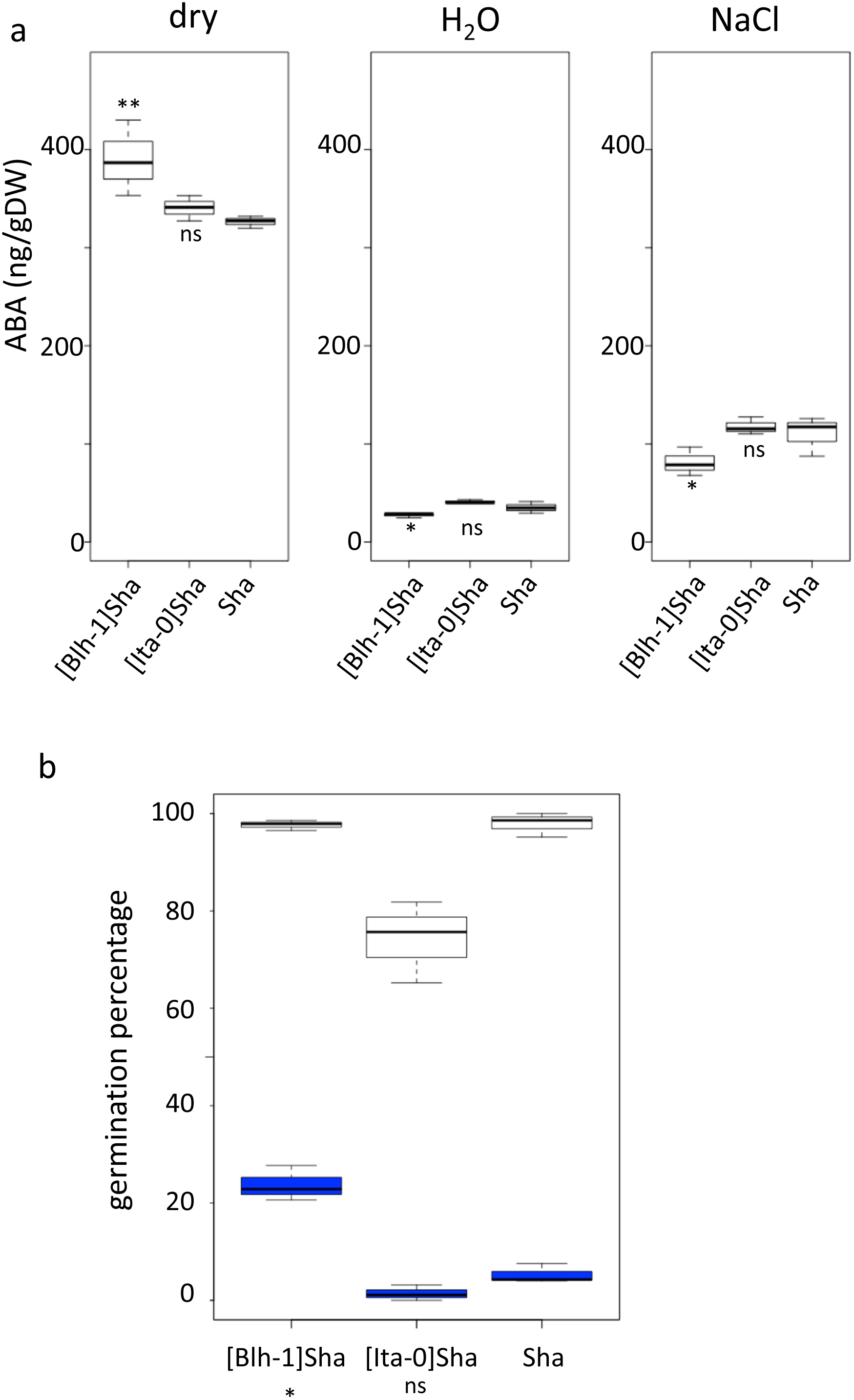
ABA content and sensitivity in seeds from lines of the Sha nucleus series. a. ABA content of seeds from [Blh-1]Sha, [Ita-0]Sha and Sha genotypes, at the dry stage (left panel), or after stratification followed by 6 hours of germination on water (middle panel) or on salt (right panel). Results of comparisons between the cytolines and Sha in the plotted experiment are shown, **<0.01<*<0.05; ns, not significant. b. Germination of stratified seeds from [Blh-1]Sha, [Ita-0]Sha and Sha on water (white boxplots) or on 50 μM ABA (blue boxplots). Germination rate was monitored after 4 days. Results of the test for a different germination response to treatment of each cytoline compared to the parent (significant cytoplasm x treatment effect) are indicated; * pval <0.05; ns, not significant (see Methods S1 for details).

## Discussion

In this study, we examined seed physiological traits important for adaptation and of relevance to agriculture, in relation to cytoplasmic genetic variants and to the coadaptation between genomic compartments. Cytolines are original and valuable genetic resources to enlighten cytoplasmic and cytonuclear effects (Roux *et al*., 2016). The analysis of a cytoline series from a given nuclear parent allows assessment of the consequences of a disruption of cytonuclear coadaptation in the considered nuclear background. The di-allele cross design used to produce the cytolines is suited to the detection of cytonuclear genetic interactions. When examining seed traits, this approach does not distinguish effects acting at the level of the mother plant from those acting at the level of the zygote(s). Nevertheless, as both the mother plant and progeny have the same genotype, it does not preclude the detection of cytoplasmic or cytonuclear effects, keeping in mind that they could act in the mother plant and/or in the progeny.

Dormancy was assessed by monitoring germination of freshly harvested seeds, or seeds after three, six or nine months of after-ripening, without stratification. Disruption of cytonuclear coadaptation had variable effects on dormancy depending on the nuclear background, both in the number of cytolines with modified dormancy and in the direction of the effect. When dormancy differed in a cytoline compared to its parent, it tended to be deeper when the parent was weakly dormant (*e.g*. Jea, Sha) and weaker in the deeply dormant Cvi-0 background. This suggests that coadapted variants of nuclear and organellar genes contributed to adaptive dormancy. However, this was not always verified: in the weakly dormant Bur-0, novel cytonuclear combinations enhanced germination. The Bur-0 nuclear series was remarkably different from the others regarding germination at the two tested temperature conditions: in contrast to the other nuclear series, the seeds of the Bur-0 nuclear series had no dormancy phenotype at 25°C (Fig. S1) but displayed dormancy at 15°C (Fig.1). A peculiar response to sowing temperature of Bur-0 seeds was also reported in a study examining a set of 73 accessions where they displayed almost 100% germination at 26°C and around 50% germination at 18°C (Schmuths *et al*., 2006). Genetic variation for the rate of dormancy loss after-ripening was previously reported for Arabidopsis (Barua *et al*., 2012). Here, by revealing that novel genetic combinations modify the kinetics of dormancy release, we provide evidence that this rate can be modulated by cytonuclear genetic interactions.

Germination performance was estimated through the germination percentage of seeds in normal and challenging conditions. The effects of foreign cytoplasms tended to lower germination performance, which is expected if optimal germinative success is assumed for natural coadapted genotypes. However, some novel cytonuclear combinations had better germination performance than their nuclear parent, suggesting that natural accessions do not always have optimal germination performance, at least in laboratory conditions. A good germination efficiency under challenging stress conditions was reported as a component of seed vigor and related to germination speed (Finch-Savage and Bassel, 2016; Yuan *et al*., 2016a). Although the limited number of time points precluded the precise fitting of germination curves and reliable comparison of germination speeds, examination of the germination time plots (Fig. S2) strongly suggested that cytolines with higher germination performance also germinated quicker than their natural parent, whereas those performing worse germinated slower. In addition, the germination percentage was influenced by cytonuclear interactions independently of the presence of salt (no third-order level of interaction), which was consistent with the idea that the detected cytonuclear effects affect germination performance per se rather than salt tolerance. This was further verified in [Blh-1]Sha and [Ita]Sha where a marker of salt tolerance (lower Na+/K+ ratio) was not linked to germination performance, whereas the specific effect of the Blh-1 cytoplasm on ABA decay rate was observed both on water and NaCl. We therefore conclude that our analysis identified effects of cytoplasm variation and cytonuclear interactions on a component of seed vigor. As longevity is also associated to seed vigor, we assessed germination after CDT in Ct-1 and Sha nuclear series, which presented contrasted sensitivities to cytoplasmic change for germination performance and dormancy. Strikingly, disruption of cytonuclear co-adaptation had no or positive effect on seed longevity, at least for the genetic variation tested. This was unexpected since lower seed longevity is generally indicative of low seed vigor and an unfit genotype. It is interesting to note that [Sha]Ct-1 and [Blh-1]Sha displayed both higher longevity and germination performance compared to their natural parents. It is conceivable that both phenotypes revealed the same underlying beneficial effect on seed vigor of the new cytonuclear combinations. However, the [Ita-0]Sha cytoline, which had a higher seed longevity than Sha, did not germinated better. A negative correlation between dormancy and seed longevity determined by co-located QTLs was observed in recombinant inbred-line populations of Arabidopsis (Nguyen *et al*., 2012). This negative correlation was not verified here: we observed deeper dormancy at 0PH and higher tolerance to CDT in the [Ita-0]Sha cytoline than in Sha. This suggests that the cytonuclear interaction effects on seed longevity and dormancy in this cytoline are not driven by the loci reported in Nguyen’s work or, not exclusively, that this cytoplasm change has different impacts on the Sha allelic effects on seed dormancy and longevity at these loci. It is also conceivable that the apparently contradiction between our result and that from Nguyen *et al*. (2012) come from differences in the conditions used to estimate seed longevity, *i.e*. survival after several years of dry storage versus CDT for 10-40 days.

Some accessions contributed more than others to the cytonuclear interacting combinations affecting germination, and individual accessions contributed differently to interactions that affect dormancy depth and germination performance after storage. For example, the Jea nucleus had the highest contribution to significant cytonuclear interacting combinations for dormancy. This suggests that, when changing the organellar partners of cytonuclear epistasis involved in dormancy tuning, Jea alleles lead to different consequences on the phenotype than alleles from other parents.

We found that the majority of cytonuclear interacting combinations that modulated germination performance also altered dormancy, with a negative correlation between dormancy and germination performance in most cases (Table 3). This is consistent with the idea that dormancy and germination speed are parts of a continuum that control germination (Finch-Savage and Leubner-Metzger, 2006). We exclude that residual dormancy of Ct-1 and Cvi-0 series induced a bias toward this conclusion because the contributions of these nuclei to significant cytonuclear interacting combinations shared by germination performance and dormancy experiments are among the fewest observed (Fig. S3). The continuity of the biological process underlying dormancy and seed vigor was recently brought back to light by a report that linked genetic variation in ABA sensitivity with seed germination performance in Brassica oleracea (Awan *et al*., 2018). ABA content was also reported to be related to germination speed in B. oleracea (Morris *et al*., 2016). In this context, our results on the [Blh-1]Sha cytoline seem particularly relevant. At the moment, a formal causal link cannot be made between the enhanced germination vigor of the cytoline on one hand, and faster ABA decay or reduced ABA sensitivity on the other hand. Nevertheless, these observations provide an interesting path for further research. In parallel to deciphering the physiological consequences of novel genetic combinations that modulate seed performance, genetic studies should also lead to the identification of gene variants that underlie the effects reported here.

One remarkable and unexpected result of the present study is the enhanced seed behavior of some cytolines compared to natural genotypes that were shaped by coevolution. Interestingly, the cytoplasm of the drought adapted Kas-1 accession was reported to have a strong and negative effect on water use efficiency in a QTL analysis (Mckay *et al*., 2008), also suggesting that the natural genotype was not optimal for this trait. It is conceivable that trade-offs between adaptive characters have led to the maintenance, in natural variants of Arabidopsis, of suboptimal genomic combinations for adaptive traits. Indeed, trade-offs between traits under selection, including germination, have been previously shown to underlie natural genetic variation in Arabidopsis (Donohue *et al*., 2005b; Todesco *et al*., 2010; Debieu *et al*., 2013). In light of these results, we suggest that more attention is given to cytoplasmic variation and its effect on seed vigor in the genetic resources of cultivated species.

## Supplementary data

Methods S1. Statistical models and contrast tests.

Fig. S1. Dormancy release during after ripening in cytoline series, germination at 25°C.

Fig. S2. Germination time curves of cytoline series on water and NaCl.

Fig. S3. Contribution of individual accessions to cytonuclear interacting combinations impacting dormancy and germination performance.

Fig. S4. Germination performance of [Blh-1]Sha and Sha seeds from two environments.

Fig. S5. Germination time curves of the Sha cytoline series on mannitol and KCl.

Table S1. NaCl concentrations used in the germination performance experiment

Table S2. Results of tests for significant cytonuclear interacting combinations in dormancy and germination performance experiments.

Table S3. Results of tests for significant effect of cytoplasms on germination after CDT.

## Acknowledgments

We thank Simon Law, Annie Marion-Poll and Helen North for helpful comments on the manuscript. We are grateful to Sébastien Thomine for his help in ion dosages. This work was funded by the Plant Biology and the Genetics and Plant Breeding INRA Departments (“Cytolignées pilotes”-2010 and “Cytoressources”-2011) and by the ANR BIOADAPT program (ANR-12_ADAP-0004). Institute Jean-Pierre Bourgin (IJPB) and Institute of Plant Sciences Paris-Saclay (IPS2) benefit from support from the LABEX Saclay Plant Sciences-SPS (ANR-10-LABX-0040-SPS).

## Abbreviations

CDT: controlled deterioration treatment
Gmax: maximum germination percentage
PH: post-harvest
RH: relative humidity

